# TARGETING 5-HT7 RECEPTOR WITH BIASED LIGANDS TO ALLEVIATE PAIN AND SPINAL NEUROINFLAMMATION

**DOI:** 10.64898/2025.12.10.693444

**Authors:** Fahima Madouri, Cyril Guimpied, Enora Pigeon, Nadège Hervouet-Coste, David Gosset, Pascal Auzou, Mélanie Kremer, Quentin Leboulleux, Marie-Aude Hiebel, Julie Le Bescont, Gérald Guillaumet, Franck Suzenet, Patrick Baril, Raphaël Serreau, Séverine Morisset-Lopez

**Affiliations:** Centre de Biophysique Moléculaire, UPR 4301 CNRS, Université d’Orléans, INSERM, rue Charles Sadron, CEDEX 02, 45071 Orleans, France; CHU, Centre Universitaire Hospitalier d’Orleans, 45100 Orleans, France; ComptOpt (Comportement et Optogénétique), Equipe Douleur et psychopathologie, INCI, CNRS UPR 3212, 67000 Strasbourg, France; Institut de Chimie Organique et Analytique, Université d’Orléans, UMR CNRS 7311, BP 6759, 45067 Orléans Cedex 2, France; UFR Sciences et Techniques, Université d’Orléans, Orléans, France; EPSM (Établissement Public de Santé Mentale) Georges Daumezon, 45400 Fleury les Aubrais, France

**Keywords:** serotonin 5-HT_7_ receptor, biased agonism, GPCR, inflammatory pain, neuroinflammation, neuropathic pain

## Abstract

The serotonin 5-HT_7_ receptor (5-HT_7_R), a member of the rhodopsin-like family of G protein-coupled receptors (GPCRs) is highly expressed in the central nervous system (CNS) and represents a promising target for treating CNS disorders such as sleep disturbances, migraine, neuropsychiatric conditions, and neuropathic pain. Owing to its therapeutic potential, extensive efforts have focused on developing selective 5-HT_7_R ligands. In the last decades, biased signalling has emerged as a key concept in GPCR pharmacology as biased ligands can stabilize specific active states of the receptor and trigger selective activation of downstream signaling pathways. In this context, we recently identified two biased 5-HT_7_R ligands, Serodolin and MOA51, from different chemical series. Here, we aimed to compare the pharmacological and safety profiles of these ligands and to assess their effect on pain-related behaviors and spinal neuroinflammation. In inflammatory pain models (acid acetic writhing, formalin and CFA tests), both serodolin and MOA51 effectively attenuated pain responses to a similar extend. Furthermore, in neuropathic pain models, spinal nerve injury (SNI) and Cuff model, both ligands reversed mechanical allodynia. Interestingly, unlike pregabalin, a clinically used reference drug, neither Serodolin nor MOA51 induced apparent tolerance after 10 consecutive days of administration. Treatment with these 5-HT_7_R ligands also reduced spinal microglial and attenuated neuronal hyperactivity in the spinal cord. Altogether these findings highlight the potential of 5-HT_7_R-biased ligands as promising analgesic candidates capable of modulating neuroinflammatory processes and mitigating both inflammatory and neuropathic pain.

## INTRODUCTION

Serotonin (5-hydroxytryptamine, 5-HT) discovered over 80 years ago, remains the focus of active research due to its broad therapeutic potential. Indeed, 5-HT and its associated receptors contribute widely in major organs systems functioning, not only gastrointestinal system where 95% of 5-HT is produced in enterochromaffin cells ^1^ but also cardiovascular, pulmonary and genitourinary system, as well as peripheral and central nervous system (CNS), which account for the remaining ∼5% of 5-HT production ^2^.

Among the 14 serotonin receptor subtypes discovered, the last identified member was 5-hydroxytryptamine 7 receptors (5-HT_7_Rs) which belong to the G protein-coupled receptor (GPCR) family. Remarkedly, nearly 40% of clinically used drugs act through GPCRs, underscoring their central role in pharmacology. The 5-HT_7_R is expressed both in many regions of the CNS such as spinal cord, thalamus, hypothalamus, cerebral cortex, amygdala and striatal complex ^3^. Owing to this broad distribution, the 5-HT_7_R has emerged as a promising target for the treatment of various CNS disorders, including depression, Alzheimer’s disease, schizophrenia, and pain ^4–6^.

Several studies have demonstrated a fundamental role of 5-HT_7_R in the control of pain ^7^. The receptor is expressed at multiple levels of the nociceptive pathway, including in nociceptors ^8^, spinal relay neurons ^9^, and supraspinal centers, notably the thalamus, the anterior cingulate cortex (ACC) ^10^, and the ventrolateral periaqueductal gray (vlPAG) ^11^. In a neuropathic pain model, the sciatic nerve injury, 5-HT_7_R is upregulated in the ipsilateral spinal cord and co-localized with GABAergic neurons ^12^.

The interest in 5-HT_7_R as a therapeutic target for pain also stems from evidence showing that its activation contributes to the analgesic effects of several clinically used drugs. For instance, spinal 5-HT_7_Rs and descending serotonergic pathways may mediate the central antinociceptive and antihyperalgesic effects of systemic paracetamol ^13^. Activation of 5-HT_7_R has also been implicated in the analgesic actions of morphine in the tail suspension test^14^ and 5-HT_7_R agonists have been shown to potentiate morphine-induced analgesia ^15^. In addition, other studies on nefopam, a widely used non-opioid analgesic in European countries for postoperative pain management ^16^, have revealed a role of the 5-HT_7_R in its antiallodynic action ^17^. Similar involvement has been reported for tramadol ^18^ and certain antidepressants such as tianeptine in a spinal cord injury models ^19^. Collectively, these results highlight the relevance of 5-HT_7_R activation in pain modulation and support the therapeutic potential of selective 5-HT_7_R agonists.

Despite these promising findings, existing 5-HT_7_R agonists suffer from important limitations, including insufficient selectivity and poor bioavailability^20^. To address these challenges, we initiated a program to identify new 5-HT_7_R ligands and subsequently discovered two compounds, Serodolin (an arylpiperazine class of molecules) and MOA51 (a benzoylpiperidine class of molecules), belonging to two distinct chemical series ^21^. Both compounds act as β-arrestin-biased ligands of 5-HT_7_R: they behave as antagonists/inverses agonists on Gs signaling while inducing ERK activation through a β-arrestin -dependent signaling ^22^. Interestingly, we also reported that Serodolin alleviates several pain-related behaviors, including mechanical allodynia and thermal hyperalgesia, in various peripheral inflammatory pain models ^22^.

In the present study, we further investigated the analgesic potential of Serodolin and MOA51 by extending our investigation to multiple inflammation and neuropathic pain murine models. Neuropathic pain does not respond to conventional pain treatments, including first-line analgesics or non-steroidal anti-inflammatory drugs (NSAIDs), making its management particularly challenging. Neuropathic pain results from damage or dysfunction of the somatosensory nervous system and is often associated with persistent symptoms such as allodynia and hyperalgesia ^23^.

A growing body of evidence indicates that neuroinflammation plays a central role in the initiation and maintenance of neuropathic pain ^24^. Following nerve injury, glial cells, especially microglia and astrocytes, are activated within the spinal cord and brain. Activated glia release pro-inflammatory mediators such as interleukin-1β (IL-1β), tumor necrosis factor-α (TNF-α), interleukin-6 (IL-6) or prostaglandins which increase neuronal excitability and contribute to central sensitization ^25^. Microglial activation occurs rapidly after injury and contributes to the early phase of pain sensitization, while astrocytes become involved in later phases, promoting the persistence of chronic pain ^26^..

In this context, our study sought to examine the therapeutic potential of two recently characterized biased 5-HT_7_R ligands, the Serodolin and MOA51, in both inflammatory and neuropathic models, while further assessing their pharmacological and safety profile. In addition, we explored their effects on molecular markers of inflammation and glial activation to better elucidate their mechanisms of action.

## RESULTS

### *In vitro* characterization of Serodolin and MOA51, molecules targeting the 5-HT_7_ serotonin receptor

The CHO-K1 cell line stably expressing the 5-HT_7A_ receptor was used to assess the specificity of Serodolin and MOA51 *(Figure 1A)* in comparison with other receptors (5-HT_1A_, 5-HT_2A_, 5-HT_2C_, 5-HT_6_, 5-HT_7A_ or D_2L_). Radioligand binding competition assays were performed to evaluate the affinity of Serodolin and MOA51 for the 5-HT_7A_ receptor. Membrane preparations expressing 5-HT_7A_R were incubated with a fixed concentration of radiolabeled ligand in the presence of increasing concentrations (10⁻¹¹ to 10⁻⁵ M) of the test compounds. 5-Carboxamidotryptamine (5-CT), a known 5-HT_7A_R agonist, was used as a reference compound *(Figure 1B)*. Serodolin and MOA51 displaced 50% of radioligand binding at nanomolar concentrations, similar to 5-CT. According to the data presented in *Table 1*, the half-maximal inhibitory concentration (IC_50_) values for Serodolin and MOA51 were 0,4 nM and 1,5 nM respectively, corresponding inhibition constants (Ki) of 0,3 nM and 1,0 nM. We further evaluated the potential off-target binding of Serodolin and MOA51 to other serotonin receptor subtypes: 5-HT_1A_R *(Figure 1C)*, 5-HT_2A_R *(Figure 1D)*, 5-HT_2C_R *(Figure 1E)* and 5-HT_6_R *(Figure 1F)*. The reference molecule for 5-HT_1A_R, 5-HT_2A_R and 5-HT_2C_R was their natural ligand, serotonin and the reference molecule for 5-HT_6_R was Mianserin, an antagonist/inverse agonist. Serodolin and MOA51 did not show strong affinity for 5-HT_1A_R *(Figure 1C)*, 5-HT_2C_R *(Figure 1E)* and 5-HT_6_R *(Figure 1F)*. Indeed, micromolar concentration are required to displace 50 % of radioligand binding. In agreement with this, IC_50_ and Ki values *Table 1* were significantly higher than those observed for 5-HT_7A_R. However, Serodolin and MOA51 also exhibited affinity for 5-HT_2A_R with Ki values of 15.4 and 2.2nM, respectively, indicating a moderate and low selectivity of 50- and 2-fold for 5-HT_7A_ versus 5-HT_2A_. Finally, when we tested binding of Serodolin and MOA51 on the dopamine D_2L_ receptor (D_2L_R) *(Figure 1G)* we found a low affinity for D_2L_R in comparison to the reference ligand risperidone. Collectively, these results indicate that both Serodolin and MOA51 are high-affinity ligands for 5-HT_7A_R, with modest affinity for other serotonin receptors expect for the 5-HT_2A_R, and minimal interaction with other evaluated dopamine receptors such as D_2L_R.

**Figure 1:**
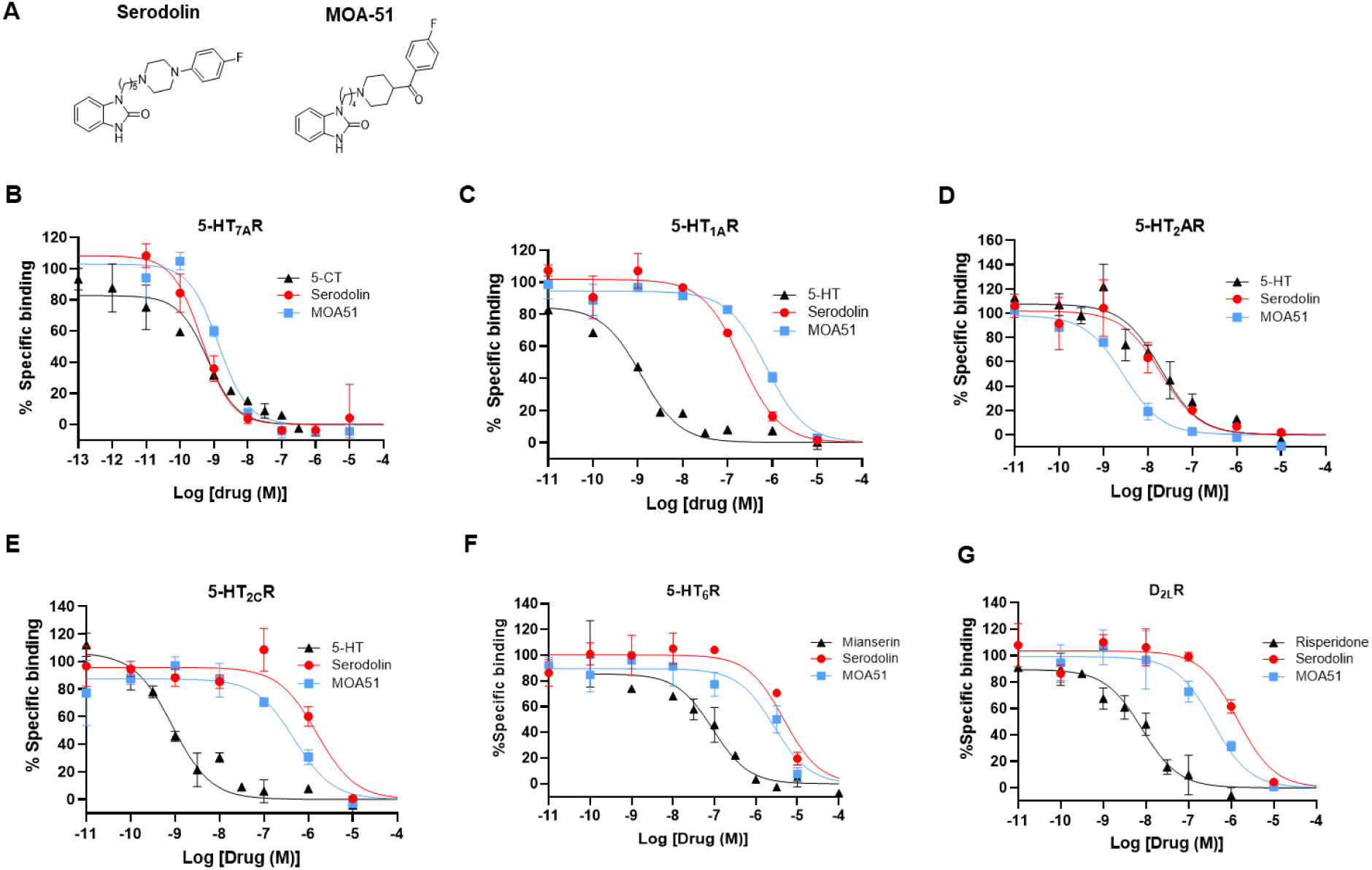
In vitro binding characterization of Serodolin and MOA51 compared to reference ligands. (A) Chemical structure of 5-HT_7_R ligands. Serodolin and MOA51.5-HT_7_R ligands were tested by radioligand binding competition assay to different receptors at seven concentrations (10^-11^M, 10^-10^M, 10^-9^M, 10^-8^M, 10^-7^M, 10^-6^M, and 10^-5^M) in duplicate. Radioligand competition assay percentage binding of ligands with 5-HT_7A_R (B), 5-HT_1A_R (C), 5-HT_2A_R (D), 5-HT_2C_R (E), 5-HT_6_R (F), Dopamine D_2L_ Receptor (G). Data are plotted from two independent experiments performed in duplicate and analysed using GraphPad prism software.

**Table 1:**
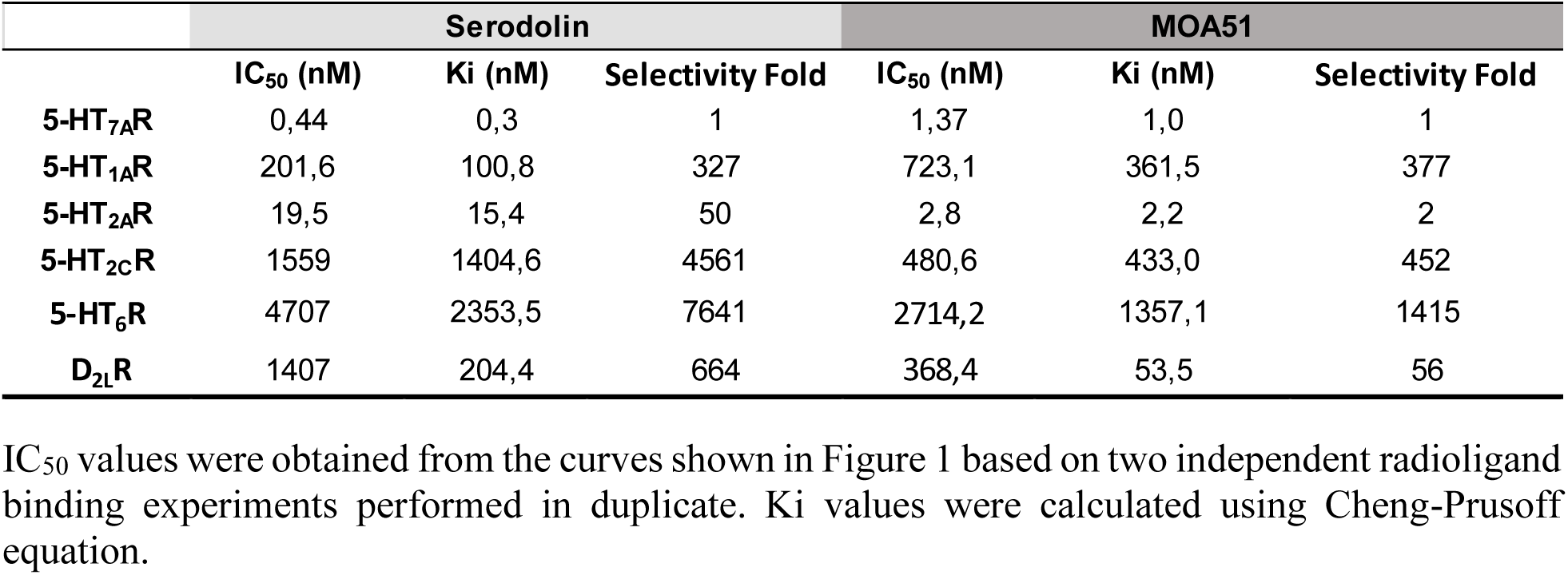
Binding affinity data of Serodolin and MOA51 for the indicated receptors.

### Serodolin and MOA51 dose-dependently reversed writhing in a visceral pain model

Analgesic activity of both compounds was first assessed using the acetic acid-induced abdominal constriction test (writhing test), a well-established model of visceral pain *(Figure 2A)*. We investigated the dose-dependent effects of Serodolin and MOA51 following a single oral administration, one hour prior to acetic acid injection *(Figure 2B-2C)*.

**Figure 2:**
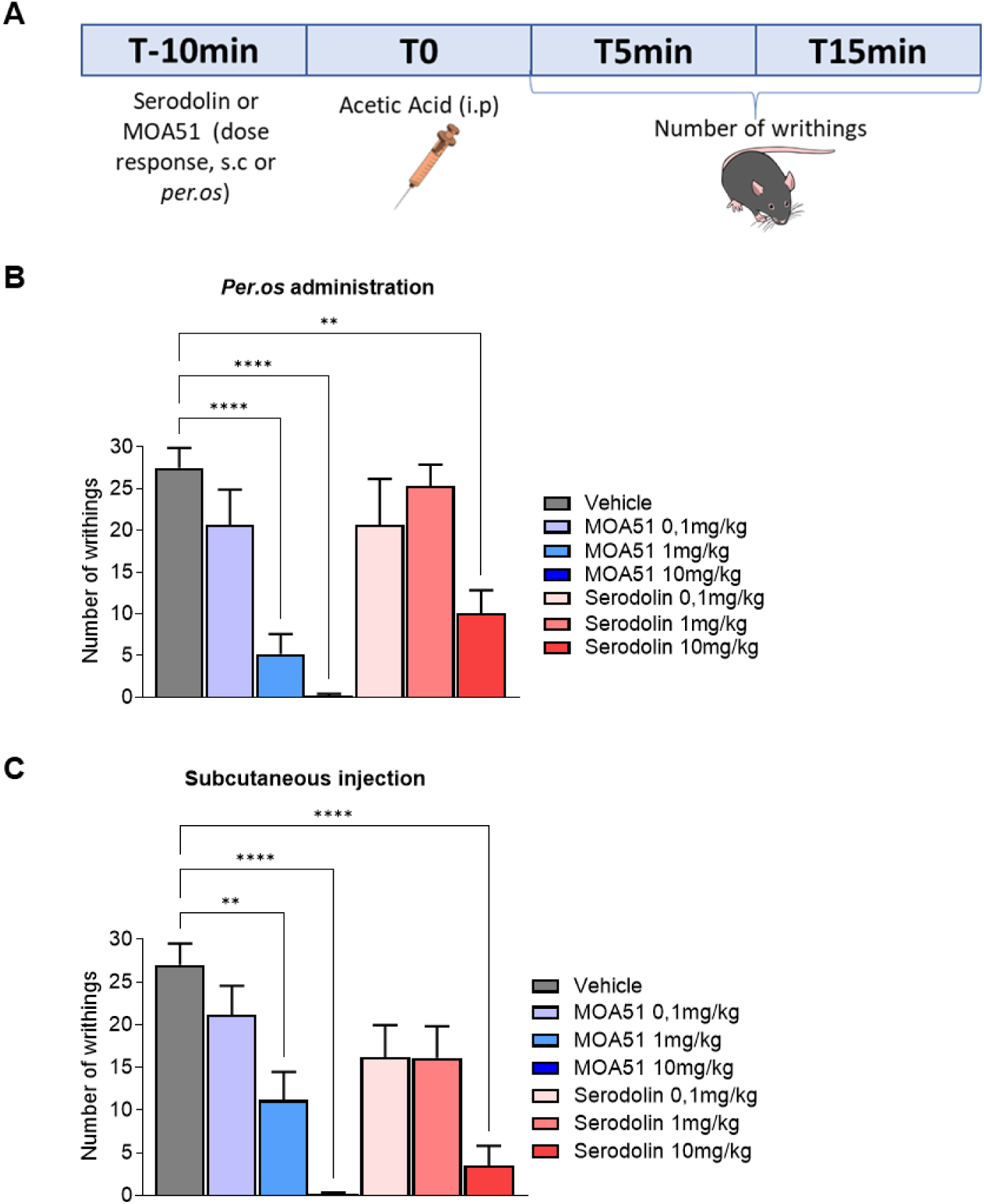
Serodolin and MOA51 reverse chronic inflammatory pain induced by acetic acid. **(A)** Design of the chronic inflammatory pain mice model induced by acetic acid. Serodolin or MOA51 were administrated at different doses (0,1 mg/kg, 1 mg/kg or 10mg/kg) per.os **(B)** or subcutaneous **(C)** route 10 minutes before the administration of Acetic Acid (i.p). Number of writhings was evaluated 5 minutes and 15 minutes after Acetic Acid for both administration routes of Serodolin an MOA51. Data are means ± SEM of values (n = 10 male Swiss mice per group). ****P < 0.0001, **P < 0.01. Statistical analysis was done using the Kruskal–Wallis test followed by non-parametric Mann-Whitney U test as compared to the vehicle-treated group.

Pretreatment with Serodolin or MOA51 resulted in a dose-dependent reduction in writhing behavior, with significant effects detected at 10 mg/kg and 1 mg/kg for Serodolin and MOA51 respectively. Therefore, we selected these two doses for subsequent experiments involving oral administration, and used half of the dose when the compounds were administered subcutaneously.

### Acute and subchronic oral administrations of Serodolin and MOA51 do not induce toxicity or blood inflammation in vivo

Before evaluating the therapeutic potential of Serodolin and MOA51 in other preclinical pain models, we aimed to verify that Serodolin and MOA51 did not induce blood toxicity or trigger inflammation associated with altered immune cells recruitment in mice. To assess the absence of toxicity of our ligands, both acute (1 day) and chronic (10 days) oral administration of vehicle (20 % DMSO, 5 % Tween 20 diluted in NaCl 0, 9%), Serodolin (10 mg/kg) or MOA51 (1 mg/kg) were performed. Twenty-four hours after the first administration, a small volume of blood was collected from the submandibular (facial) vein (day 1). For chronic treatment, blood was collected intracardially at day 10. Compared to the Vehicle-treated group, our results demonstrate that Serodolin and MOA51 did not cause any significant increase or decrease in the number of white blood cells *(Figure S1A)*, red blood cells (Erythrocytes) *(Figure S1B)*, mean platelet volume *(Figure S1C)*, hemoglobin *(Figure S1D)* and the percentage of hematocrit (Figure S1E). The blood cell counts remained unchanged following administration of either Serodolin or MOA51. Taken together, these results show that both acute and chronic oral administration of Serodolin and MOA51 did not induce anaemia or erythrocytosis, nor did not lead to leukopenia or leucocytosis. Moreover, we assessed the proportions of immune cell populations in peripheral blood at day 1 and day 10 post-treatment. Specifically, we analysed lymphocytes *(Figure S1F)*, granulocytes *(Figure S1G)*, eosinophils *(Figure S1H)* and monocytes *(Figure S1I)*, which are typically elevated during systemic inflammation. Serodolin and MOA51, whether administered acutely and chronically did not alter the percentages of these immune cell subsets in comparison to vehicle-treated groups. We also measured platelet (thrombocytes) counts *(Figure S1J)*, which are critical for coagulation, and found no difference among groups. Additionally, body weight of animal was also monitored throughout the chronic treatment period for all groups. No significant weight loss was observed in mice treated with Serodolin or MOA51 compared with vehicle-treated controls *(Figure S1K-L)*. Altogether, our data indicate that both acute and chronic oral administration of Serodolin and MOA51 did not induce haematological toxicity or systemic inflammation and had no effect on immune cell recruitment in the blood of treated mice.

### Serodolin and MOA51 reverse CFA-induced mechanical allodynia in WT mice but not in 5-HT_7_R knockout mice

In our previous study ^22^, we explored the antinociceptive activity of Serodolin *in vivo* following complete Freund’s adjuvant (CFA) sensitization. We showed that mice injected with Serodolin displayed a reduction of the mechanical threshold triggering withdrawal of the ipsilateral paw in the Von Frey test. Moreover, pre-treatment of animals with SB269-970, a 5-HT_7_R antagonist, 30 min before Serodolin administration fully blocked its effect on allodynia. Thus, in the present study, we wanted to go deeper in this and explore the effect of both Serodolin and MOA51 on mechanical allodynia induced by CFA in Wild Type (WT) and also in 5-HT_7_R knock out mice (5-HT_7_R KO mice). As described in *Figure 3A*, the mechanical sensitivity of the mice was assessed using Von Frey filament at day 0 to determine the baseline response. On day 1 we evaluated the effect of CFA. Both WT and 5-HT_7_R KO mice injected with CFA into the midplantar surface of the left hindpaw (ipsilateral paw) developed a significative mechanical hypersensitivity in comparison to the baseline. Subcutaneous injection of Serodolin (5 mg/kg) or MOA51 (0,5 mg/kg) reversed the CFA-induced mechanical hypersensitivity in WT mice *(Figure 3B-C)*. Importantly, CFA-induced mechanical hypersensitivity was not reversed when Serodolin or MOA51 were injected to 5-HT_7_R KO mice *(Figure 3D-E)*. Thus, these data suggest that the anti-allodynic effect of Serodolin and MOA51 observed in CFA-induced mechanical hypersensitivity was triggered by 5-HT_7_R.

**Figure 3:**
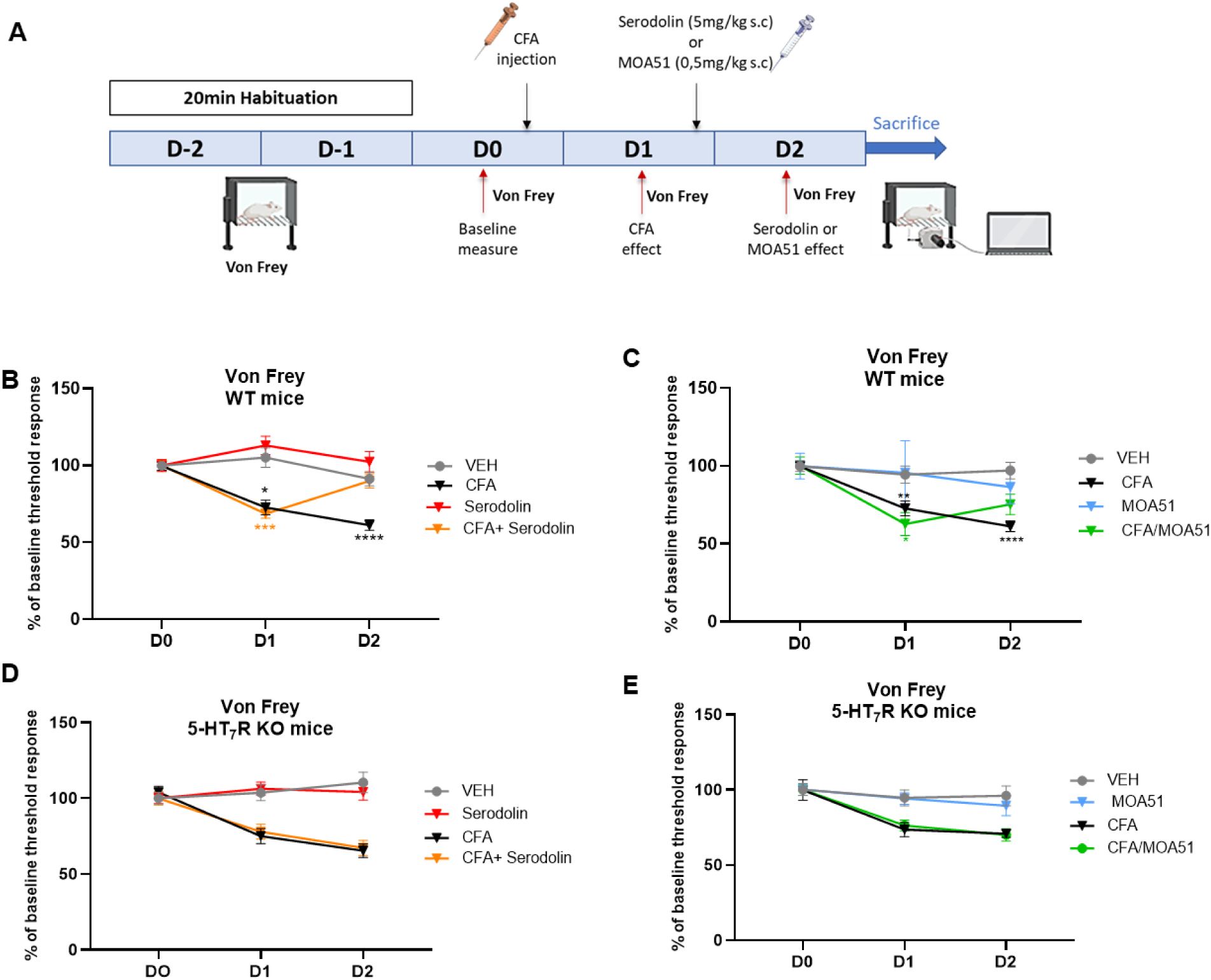
Serodolin and MOA51 reverse CFA-induced mechanical allodynia in WT mice but not in 5-HT_7_R knockout mice. **(A)** Design of the mechanical allodynia induced by Complete Freund Adjuvant (CFA). After a habituation period in the apparatus, baseline Von Frey measures were realized on day 0, then CFA injection in the left hind paw was performed. The effect of CFA was evaluated on mechanical allodynia with Von Frey apparatus 24h after CFA injection on day 1. Serodolin (5 mg/kg, s.c) **(B)** or MOA51 (0,5 mg/kg, s.c) **(C)** were administrated on day 1 and their effect on mechanical allodynia was evaluated with Von Frey 24h later, on day 2. Von Frey measurement on 5-HT_7_R knock out (KO) mice following injection of CFA and Serodolin (5 mg/kg, s.c) **(D**) or MOA51 (0,5 mg/kg, s.c) **(E)**. Results were normalized as the percentage of baseline threshold response. Data are means ± SEM of values obtained in two independent experiments (n=5 to 10 mice per group, male and females). Statistical analysis were realized with comparison to the D0 respectively for each condition. ***P < 0.001 Statistical analysis was done using the Tukey’s test after significant two-way repeated measures ANOVA.

### Serodolin and MOA51 exhibits anti-inflammatory effects comparable of morphine in a rat model of acute pain induced by formalin injection

We evaluated the anti-inflammatory effect of Serodolin and MOA51 in rats using the formalin test. As shown in *Figure 4A*, Serodolin (10 mg/kg), MOA51 (1 mg/kg) or Morphine (1 mg/kg) were subcutaneously injected in rats. Then 30 min after the injection, a 2,5 % formalin solution was administrated into the plantar aspect of the hindpaw. This solution produces a biphasic painful response. The early phase of the response begins immediately after the formalin injection and it last 5 minutes. This initial phase was caused by direct activation of peripheral nociceptors by formalin. The late phase of the response was evaluated between 17 and 27 minutes and results from inflammatory process and central sensitization in the spinal cord.

**Figure 4:**
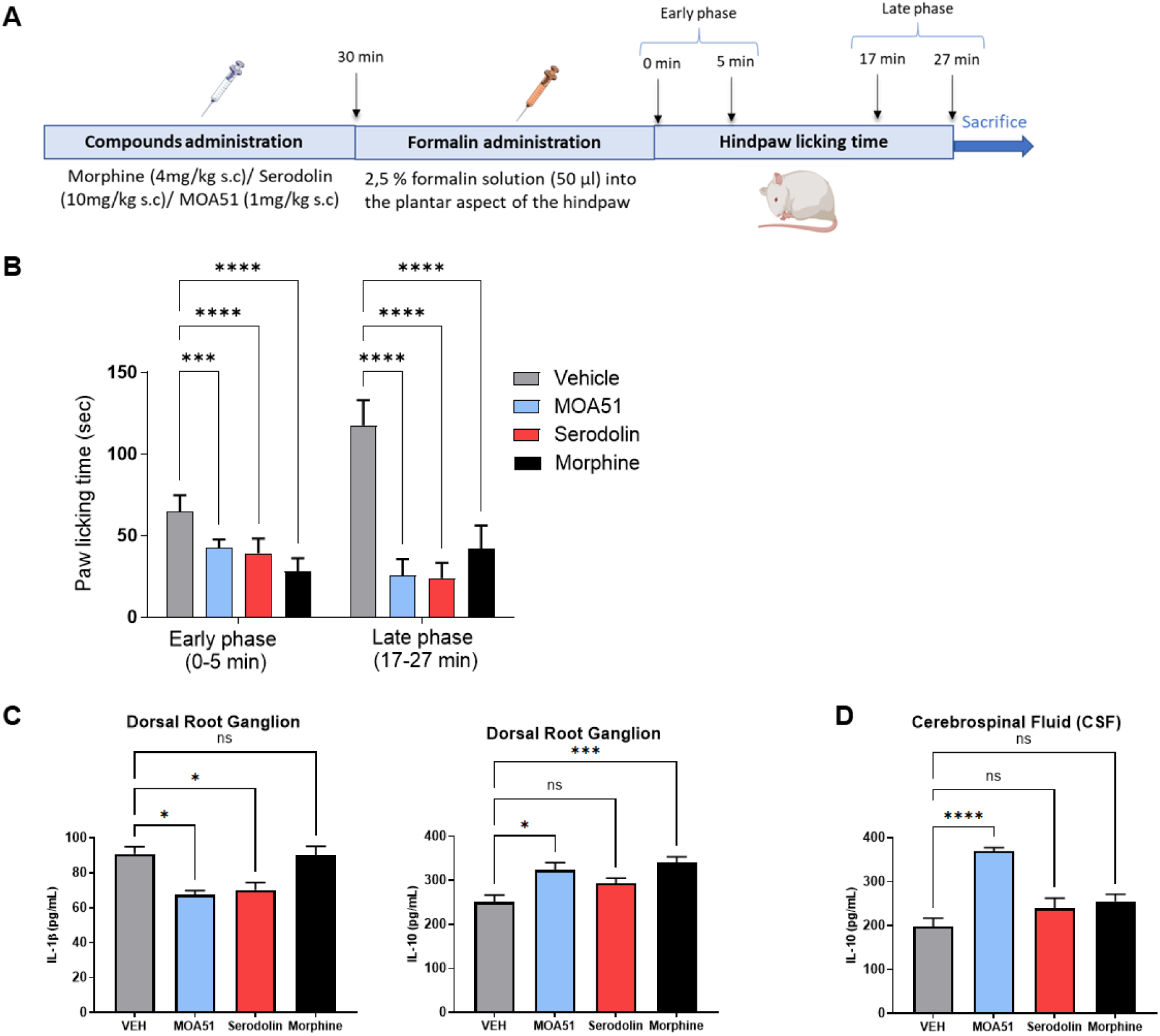
Serodolin and MOA51 produced anti-inflammatory effects comparable to those of morphine in an acute pain rat model (formalin test). **(A)** Design of Formalin rat model experiment. **(B)** Paw licking time was measured from 0 to 5 min after formalin injection (early phase) and from 17 to 27 min after formalin injection (late phase). Morphine (4 mg/kg, s.c), Serodolin (10 mg/kg, s.c) or MOA51 (1 mg/kg, s.c) were administrated 30 min before injection of 2,5 % formalin solution into the plantar aspect of the hindpaw. **(C)** IL-1β (pg/mL) and IL-10 (pg/mL) production by right Dorsal Root Ganglia (DRG) was measured by Enzyme-Linked Immunosorbent Assay (ELISA). **(D)** IL-10 (pg/mL) production in Cerebrospinal fluid was measured by ELISA. Data are means ± SEM of values (n = 8 male Sprague-Dawley rats per group). ****P < 0.0001, ***P < 0.001, *P < 0.05. Statistical analysis was done using the Kruskal–Wallis test as compared to the vehicle-treated group.

Both MOA51 and Serodolin significantly reduced paw licking time during both the early and late phase, compared to vehicle-treated rats *(Figure 4B)*. These results demonstrate the promising analgesic efficacy of Serodolin and MOA51. At the end of behavioural experiments, dorsal root ganglion (DRG) from lumbar 4 to 6 were collected as well as cerebrospinal fluid. We evaluated by Enzyme-Like Immunosorbent Assay (ELISA), the production of a pro-inflammatory cytokine (IL-1β) and an anti-inflammatory cytokine (IL-10) by DRG *(Figure 4C)*. Interestingly, IL-1β production was significantly reduced in dorsal root ganglion of rats treated with Serodolin and MOA51 but not those treated with Morphine. Moreover, treatment with MOA51 and Morphine led to a significant increase in IL-10 levels, an effect not observe with Serodolin-treated rat. When we assessed the levels of IL-1β levels in the cerebrospinal fluid (CSF), we were unable to detect its presence, most likely due to the detection limit of the assay. However, we found a significant increase of IL-10 production in CSF of rats treated with MOA51, but not those treated with Serodolin or morphine *(Figure 4D)*. Overall, our results strongly suggest that both Serodolin and MOA51 exhibit analgesic and anti-inflammatory effect comparable to those of morphine in an acute formalin-induced pain model in rats. Notably, MOA51 showed a particularly protective effect by increasing IL-10 production, an anti-inflammatory cytokine, in both the DRG and CSF.

### Serodolin and MOA51 *per.os* administration have a complete and satisfactory effect on mechanical allodynia in the so-called “Cuff” model of sciatic nerve injury

After demonstrating the anti-allodynic, analgesic and anti-inflammatory effects of Serodolin and MOA51, we explored their potential impact on neuropathic pain using the “cuff” model of sciatic nerve injury *(Figure 5A)*. The neuropathic pain was induced by compression of the main branch of the right sciatic nerve. Animals with surgery were part of the “Cuff” group. The “Sham” control animals underwent the same surgery without cuff insertion. Duloxetine (30 mg/kg) was used as a reference molecule and different doses of Serodolin (0,1 mg/kg, 1 mg/kg, 5 mg/kg, 10 mg/Kg) or MOA51 (0,01 mg/kg, 0,1 mg/kg, 0,5 mg/kg, 1 mg/kg) were orally administered 15 days after the surgical procedure in order to defined the optimal concentration. Sham surgery, as a negative control, did not influence mechanical thresholds as compared with baseline measurement. Serodolin at 10 mg/kg *(Figure 5B)* and MOA51 at 1 mg/kg *(Figure 5C)* showed a complete and satisfactory effect on mechanical allodynia in the “cuff” model of sciatic nerve injury outperforming Duloxetine, the reference molecule, which was administrated at a higher dose. The other doses tested did not shown an anti-allodynic effect, so we therefore used the optimal doses to performed ELISA on the plasma samples of mice treated with the effective doses of Serodolin (10 mg/kg) and MOA51 (1 mg/kg) *(Figure 5D)*. Interestingly, we observed a significant decrease of IL-1β production in plasma samples of mice treated with 10 mg/kg of Serodolin compared to cuff vehicle mice. Although not significant, both MOA51 and Duloxetine also reduced IL-1β amount in plasma. Moreover, we evaluated the production of IL-12p40, a cytokine primarily produced by activated inflammatory cells. We showed that IL-12p40 was significantly reduced in plasma of mice treated with Serodolin and Morphine. Even if it was not significant in the plasma samples of mice treated with MOA51, there was also a decreased production of IL-12p40 compared to cuff vehicle mice. Unfortunately, we were unable to measure IL-10 levels in the plasma due to the detection limit of the assay. Finally, we evaluated the production of TNF-α, an inflammatory cytokine produced during acute inflammation. Both Serodolin and MOA51 reduced TNF-α levels in plasma samples. To reinforce our data, we conducted the same experiment but following subcutaneous injection of Serodolin (0,1 mg/kg, 1 mg/kg, 2,5 mg/kg, 5 mg/kg) or MOA51 (0,01 mg/kg, 0,1 mg/kg, 0,25 mg/kg, 0,5 mg/kg) *(Supp Data Figure S2)*. Serodolin significantly alleviated the cuff-induced allodynia at several doses (0,1 mg/Kg, 2,5 mg/kg, 5 mg/kg) while MOA51 was effective at 0,25 mg/kg and 0,5 mg/kg. Altogether, our results demonstrated the anti-allodynic and anti-inflammatory effect of our ligands in the so-called “Cuff” model of sciatic nerve injury.

**Figure 5:**
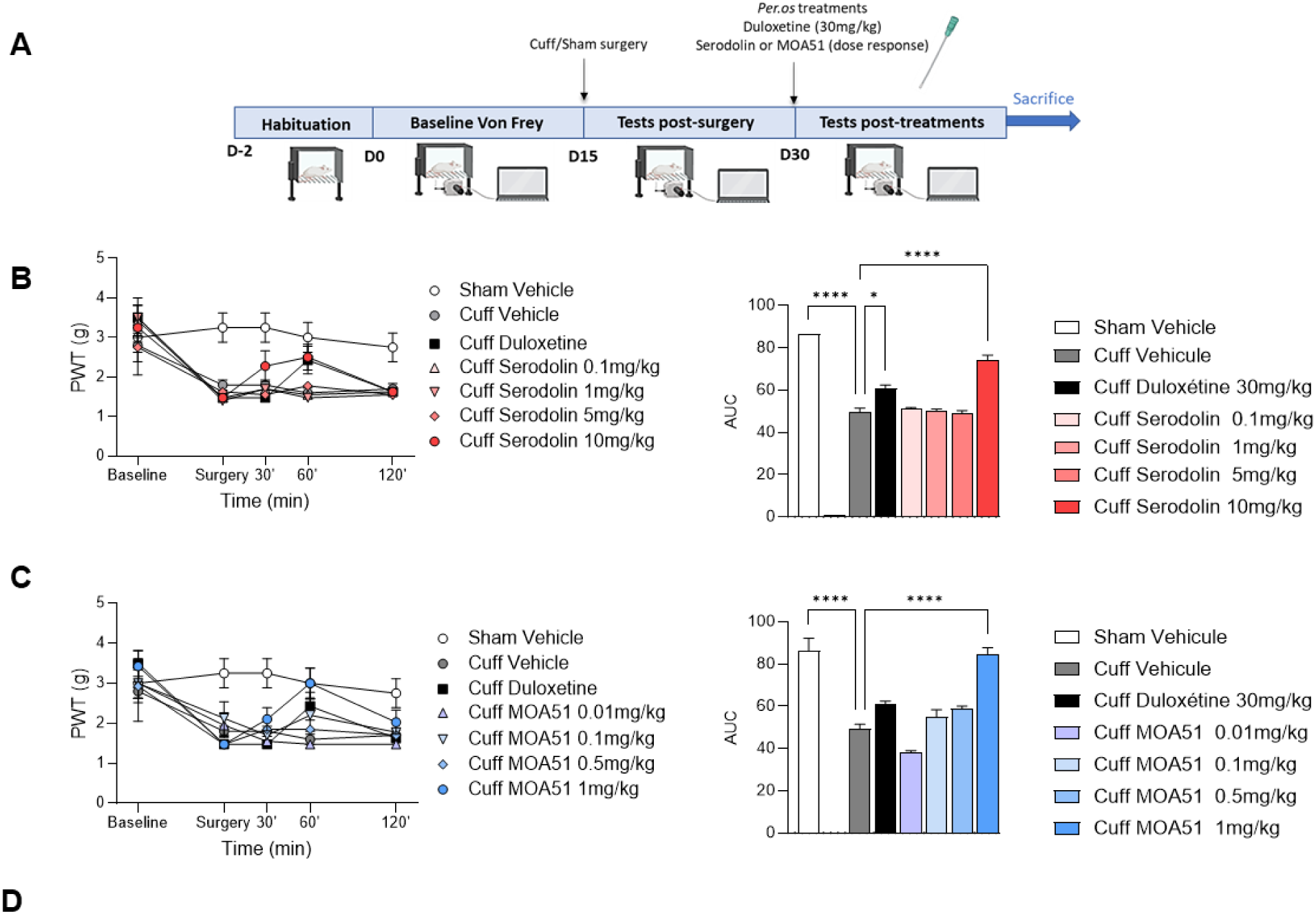
Oral administration of Serodolin and MOA51 fully alleviates mechanical allodynia in the “Cuff” model of sciatic nerve injury. **(A)** Design of “Cuff” model of sciatic nerve injury mice (n=7 mice per group). After a habituation period in the apparatus, baseline Von Frey measurements were realized on day 0. On day 15, cuff/sham surgery was performed and Von Frey measurement were realized to verify the effect of the surgery. Mice were treated with Duloxetine (30 mg/kg, s.c) or different doses of Serodolin per.os administration (10 mg/kg; 5 mg/kg; 1 mg/kg; 0,1 mg/kg) or MOA51 per.os administration (1 mg/kg; 0,5 mg/kg; 0,1 mg/kg; 0,01 mg/kg). **(B)** The effect of Serodolin was evaluated on mechanical allodynia with Von Frey. **(C)** The effect of MOA51 was evaluated on mechanical allodynia with Von Frey. Results were represented as curve with Time Course and as Area Under the Curve (AUC). The results are expressed as the group means ± standard error of the mean (SEM). The AUCs were calculated using the trapezoid method taking the value 0 as a reference. Il-1β, IL-12p40 and TNF-α production in plasma samples were evaluated using ProcartaPlex from ThermoFisher Scientific **(D)**. The analysis of the results was carried out using Statistica 13 software (Statsoft, USA). A statistical analysis was performed using a multifactorial analysis of variance (ANOVA) during repeated measurements. The surgical procedures (“Sham” or “Cuff”) and the different treatments were selected as between-group factors. The measurement time was taken as a within-subject factor. The Duncan test was used for post-hoc comparison. The statistics are represented in the results section compared to the “Cuff Vehicle” group with *p<0.05, **p<0.01, ***p<0.001 and ****p<0.0001.

### Serodolin and MOA51 reverse mechanical allodynia with no apparent tolerance when administered daily for nine consecutive days after spared nerve injury surgery

Finally, to confirm the potential of Serodolin and MOA51 to treat chronic pain, we evaluated their effects in another mice model of neuropathic pain, the Spared Nerve Injury (SNI) *(Figure 6A)*.

**Figure 6:**
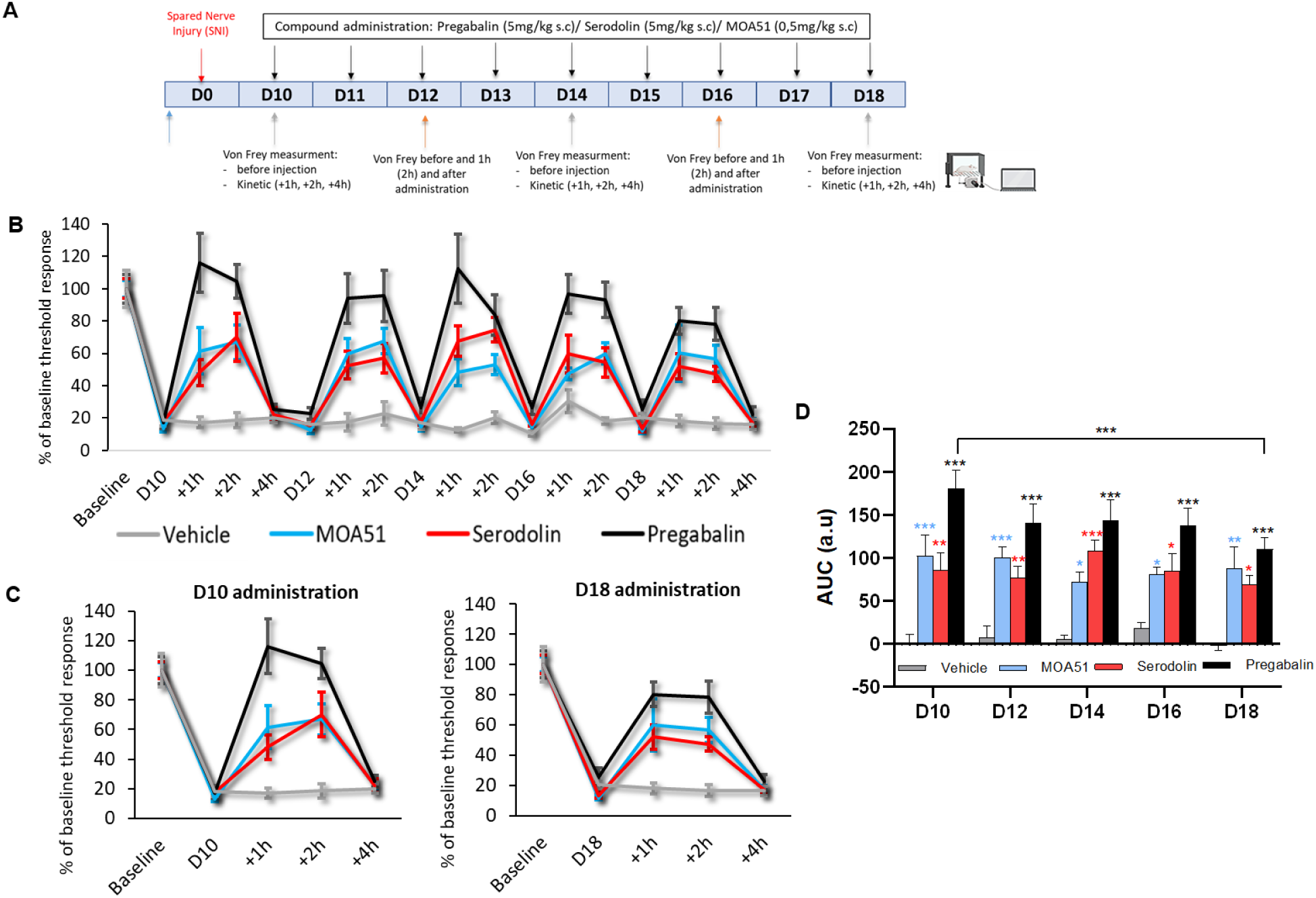
Serodolin and MOA51 reverse mechanical allodynia without apparent tolerance after daily administration for nine consecutive days following spared nerve injury surgery. **(A)** Design of Spared Nerve Injury (SNI) mice model. Surgery was done at day 0 and compound was injected subcutaneaously daily during 8 days from day 10 to day 18 (Pregabalin, 5mg/kg, Serodolin, 5mg/kg and MOA51, 0,5mg/kg). Timepoint of Von Frey measurement are indicated in the figure. **(B)** The effect of compounds was evaluated on mechanical allodynia with Von Frey. Results were represented as percentage of baseline threshold response. **(C)** The effect of compounds at day 10 (first administration) and day 18 (last administration) was evaluated on mechanical allodynia with Von Frey. Results were represented as percentage of baseline threshold response. **(D)** Area Under the Curve (AUC) of time-course for indicated days for repetitive administration of Vehicle, MOA51, Serodolin and Pregabalin in SNI neuropathic pain model. n=10 mice per group were used in this study. Two-way RM ANOVA (followed by Bonferroni post-hoc test) was used to analyse time course response day 10 and day 18. Area under the curve was determined with the 1h and 2h post-drug administration measures to investigated tolerance of repeated administration of compounds.

A complete ligature and transection of the common peroneal and tibial distal branches of the sciatic nerve was made leaving the sural branch intact. Mechanical threshold responses in mice were measured every two days with calibrated Von Frey filaments following daily subcutaneous administration of Serodolin (5 mg/kg) or MOA51 (0,5 mg/kg). Pregabalin (5 mg/kg), the reference molecule produced a strong anti-allodynic effect on spared nerve injury-induced mechanical allodynia, observable even after the first dose *(Figure 6 B-D)*. Although the effect of Serodolin and MOA51 were less pronounced than those of Pregabalin after the first administration, interestingly, they exhibited a stable effect over time (from day 10 to day 18) unlike Pregabalin, whose efficacy significantly diminished after ten repetitive injections *(Figure 6 C-D)*. Overall, our results showed that Serodolin and MOA51 reverse mechanical allodynia with no apparent tolerance when administered daily for nine consecutive days after spared nerve injury surgery.

### SNI-induced activation of astrocytes, microglia, and neurons is markedly reduced in the presence of Serodolin and MOA51

Finally, to gain insight into the molecular mechanism underlying the therapeutic effects of MOA51 and Serodolin in the SNI model of neuropathic pain, we collected spinal cords to evaluate neuroinflammation processes through immunofluorescent staining of various neurological cell types: Glial Fibrillary Acidic Protein (GFAP) for astrocytes detection *(Figure 7A)*, Iba-1 (Ionized calcium-binding adaptor molecule 1) for microglia detection *(Figure 7B)* or c-Fos (a marker of neuronal activity) *(Figure 7C)*. We observed a significant decrease in astrocytes activation in mice treated with Serodolin, MOA51 or Pregabalin, with the strongest effect seen with Serodolin *(Figure 7A)*. Moreover, microglia and neurons activation were significantly reduced only in the Serodolin-treated group, although quantifications of the staining for MOA51 or pregabalin also showed some reduction compared to Vehicle-treated SNI mice *(Figure 7B-C)*. These findings, combined with our behavioural strongly support the conclusion that Serodolin and MOA51 are effective in the treatment of neuropathic pain.

**Figure 7:**
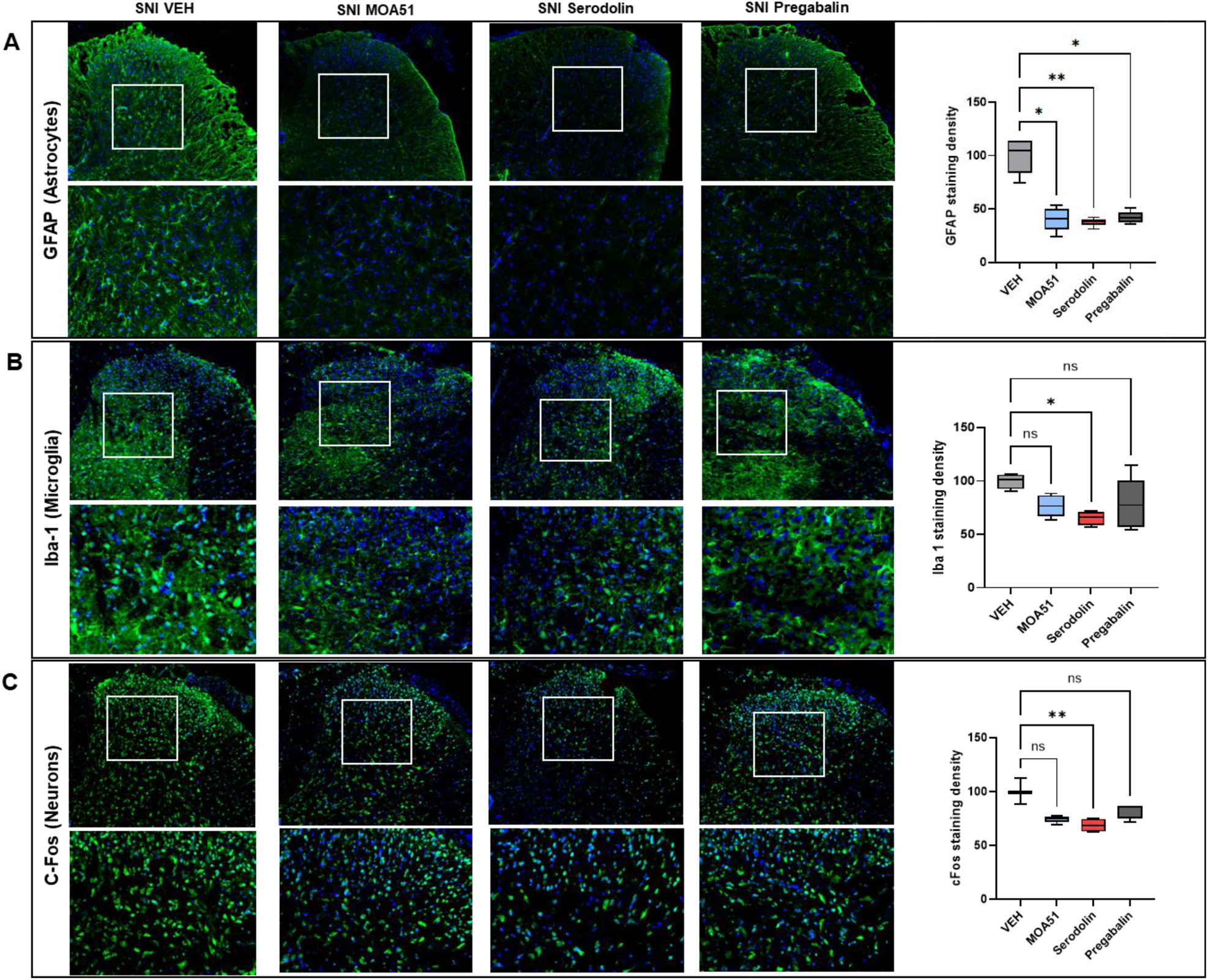
Astrocytic, microglial, and neuronal activation induced by SNI surgery is reduced by Serodolin and MOA51. **(A)** GFAP (Glial Fibrillary Acidic Protein, an astrocytic marker), **(B)** Iba-1 (Ionized calcium-binding adaptor molecule 1, a microglia marker) and **(C)** c-Fos (a marker of neuronal activity) were stained in spinal cord of mice with SNI surgery treated or not with MOA51, Serodolin or Pregabalin. Images quantification, performed on the entire slices, shows a significant decrease in GFAP staining density in the ipsilateral lumbar spinal cord of SNI mouse treated with MOA51, Serodolin and Pregabalin, a significant decrease in Iba-1 and c-Fos staining density in spinal cord of SNI mouse treated with Serodolin. Photos was taken with 20×0.8 objective. A focus was done on spinal dorsal horn. Scale bars = 50 μm. Data were means ± SEM of values. Three spinal cord per group were used for immunohistological staining. Six slices per mouse, taken between Lumbar segments (L3) and (L5) were stained with fluorescent antibodies. ****P < 0.0001, ***P < 0.001, **P < 0.01, *P < 0.05. Statistical analysis was done using the Kruskal–Wallis test as compared to the vehicle-treated group.

## DISCUSSION

Over the past two decades, several selective 5-HT_7_R ligands have been developed. However, many of these compounds display limited specificity or suboptimal pharmacokinetic properties, thereby restricting their therapeutic applicability ^27^. In our previous study, we identified two novels 5-HT_7_R-β-arrestin-biased ligands, named Serodolin (an arylpiperazine derivate) and MOA51 (a benzoylpiperidine class of molecules), belonging to distinct chemical series ^21–22^. Biased ligands can preferentially activate specific intracellular pathways, offering the possibility of enhanced efficacy and reduced side effects compared with conventional GPCRs agonists ^28–30^.

To further characterize Serodolin and MOA51, we performed radioligand binding assays across various serotonin receptor subtypes and dopamine D_2_ receptor. Serodolin exhibited high affinity for the 5-HT₇ receptor (Kᵢ = 0.3 nM) and excellent selectivity while MOA51 also bound to the 5-HT₇ receptor with high affinity (Kᵢ = 1 nM) and favorable selectivity ratios. In line with previous pharmacomodulation data ^21^, Serodolin displayed higher selectivity over 5-HT_2A_ than MOA51. Given the well-documented hallucinogenic potential associated with 5-HT₂_A_ receptor agonists ^31^, it was reassuring to observe that they act as antagonists at the 5-HT_2A_ receptor ^21^. Considering the relatively high affinity of our compounds for the 5-HT_2A_ receptor, we cannot fully exclude its involvement in pain modulation. However, results from 5-HT_7_ knockout mice demonstrate a specific action through 5-HT_7_R, with no evidence supporting a role for 5-HT_2A_. Consistently, our previous work in a CFA mouse model showed that 5-HT7R is essential for Serodolin’s anti-allodynic effects, as these effects were completely abolished in 5-HT7R knockout mice or after treatment with the selective antagonist SB-269970 ^32^.

Beyond efficacy, safety is a prerequisite for drug development. Remarkably, acute and chronic oral administration of Serodolin or MOA51 alone did not disrupt immune cell balance (lymphocytes, granulocytes, eosinophils and monocytes) and did not alter key blood parameters, including white blood cells, red blood cells, platelet, hemoglobin, hematocrit ^33^. This, indicate the good tolerability of Serodolin and MOA51 in mice in a non-pathogenic context.

Due to its localization both in the peripheral and in the central nervous system ^3^, over the last few years, there were several evidences showing the fundamental role of the 5-HT_7_R in the control of pain and its potential therapeutic applications ^7, 34^. Numerous studies have evaluated the effect of either agonists or antagonist of the 5-HT_7_R in the modulation of pain. For example, microinjection of the 5-HT_7_R agonist AS-19 into the ventrolateral midbrain periaqueductal gray (vlPAG) produced dose-dependent anti-hyperalgesia effects in the chronic constriction injury model, which were reversed by the selective 5-HT_7_R antagonist SB269970 ^35^. Similarly, targeting 5-HT_7_R with the agonist E-57431 modulated neuropathic pain in the partial sciatic nerve ligation model ^12^. Recently, we demonstrated that Serodolin showed potential analgesic in persistent pain states involving tissue damage, as observed in two inflammatory pain models: the CFA-induced inflammation model and the formalin-induced pain model ^22^. Here, both MOA51 and Serodolin significantly reduced allodynia in the CFA mice model, an effect absents in 5-HT₇R knockout mice. These results support and extend the previous data obtained in the tail immersion test, showing that the reduction in tail withdrawal latency induced by subcutaneous administration of E55888 or Serodolin was not observed in 5-HT_7_R KO mice ^22^.

The formalin-induced pain model is also a widely used model to evaluate the impact of compounds on inflammation-induced pain. It has been employed to assess the effects of 5-HT₇R ligands such as LP-44 and LP-211 ^36^ and AS-19 ^37^. By using this model, we demonstrated that Serodolin and MOA51 produced anti-inflammatory effects comparable to those of Morphine and that both ligands exhibited an analgesic effect by reducing the paw licking time. It is known that subcutaneous injection of formalin into the rat hind paws, leads to an increase in interleukin-1β (IL-1β) production, a pro-inflammatory cytokine, in the brain ^38^. We also demonstrated that Serodolin and MOA51 exhibited anti-inflammatory effects by reducing IL-1β production in dorsal root ganglion (DRG) and by increasing IL-10 levels, an anti-inflammatory cytokine, in both the DRG and cerebrospinal fluid (CSF). These cytokine changes likely contribute to their analgesic effects and underscore the potential of Serodolin and MOA51 as promising candidates for the treatment of inflammatory pain.

Neuropathic pain is caused by lesions or pathology of the somatosensory nervous system, which largely alter the quality of life of the patients ^39^ and responds poorly to most analgesics, with the exception of certain opioids ^40^. However, due to their long-term side effects, such as dependence and tolerance, opioids are generally not suitable for chronic pain management. As a result, the main treatments currently used are antidepressants and antiepileptics ^41–42^, which tend to have fewer side effects. Nonetheless, their effectiveness remains moderate and is observed in only about 50% of patients ^39^, highlighting the need for more effective therapies. To address this, we assessed Serodolin and MOA51 in two models of neuropathic pain: sciatic nerve compression (cuff model) ^43–44^ and transection (SNI model) ^45^. To assess their translational potential, we evaluated the pharmacological effects of MOA51 and Serodolin following different routes of administration (subcutaneous and oral) and at multiple doses. In the sciatic nerve cuff model, oral administration of Serodolin (10 mg/kg) and MOA51 (1 mg/kg) produced a robust and complete anti-allodynic effect and, remarkably, to a higher extent compared to reference compound Duloxetine, even while administrated at higher dose (30 mg/kg). Subcutaneous administration of Serodolin also alleviated the cuff-induced allodynia accross several doses (0,1 mg/Kg, 2,5 mg/kg, 5 mg/kg), while MOA51 was effective at 0,25 mg/kg and 0,5 mg/kg. Moreover, Serodolin decreased plasma levels of IL-1β and IL-12p40 and both Serodolin and MOA51 decreased TNF-α production. These findings suggest that, beyond alleviating neuropathic pain symptoms, both ligands also reduce systemic inflammation, which may contribute to their overall therapeutic effect. Overproduction of IL-1β in the injured sciatic nerve has been proposed as a common mechanism underlying the neuropathic pain, memory impairment and depression ^46^. TNF-α has also been strongly implicated in the physiopathology and peripheral mechanisms of neuropathic pain ^47–48^.

Interestingly, in SNI model, both compounds reversed mechanical allodynia without apparent tolerance over nine consecutive days. Pregabalin (5 mg/kg, *s.c*), used as a reference, produced significant analgesia compared to vehicle but showed a marked decline in efficacy between the first and last administration, indicating tolerance. In contrast, Serodolin (5 mg/kg, *s.c*) and MOA51 (0,5 mg/kg, *s.c*), maintained stable analgesic effects over time, slightly lower than pregabalin but without signs of tolerance. To investigate the underlying mechanisms, spinal cord tissue was analysed via immunofluorescent staining. Since 5-HT_7_R is expressed on astrocytes, microglia and neurons ^6^, we focused our investigation on astrocyte activation using Glial Fibrillary Acidic Protein (GFAP) staining ^30^, microglia with Iba-1 (Ionized calcium-binding adaptor molecule 1) ^49^ and neuronal activity with c-Fos ^50^. We found that astrocytes activation was significantly decrease in mice treated with Serodolin, MOA51 or Pregabalin, with the strongest effect observed for Serodolin. Moreover, only Serodolin significantly decreased microglial and neuronal activation, although MOA51 and pregabalin showed a tend toward reduction compared to Vehicle group. Therefore, administration of 5-HT_7_R-biased ligands effectively counteracted SNI-induced glial activation and mechanical allodynia in mice, reinforcing the contribution of glial mechanisms to the analgesic effects of 5-HT_7_R ligands. This observation is consistent with preclinical evidence showing that conventional analgesics are capable of modulating glial responses. For instance, Ammoxetine, a serotonin reuptake inhibitor, demonstrated strong analgesic efficacy and attenuate spinal glial activation and inflammatory mediator release triggered by peripheral nerve injury ^51^. Likewise, Amitriptyline reduces spinal microglial and astrocytic activation following nerve injury while producing robust antinociceptive effects ^52^. Gabapentin similarly dampens microglial activation in models of diabetic neuropathy and improves pain-related behaviors ^53^.

Altogether, our data demonstrate that the biased 5-HT_7_R ligands Serodolin and MOA51 alleviate inflammatory and neuropathic pain, reduce central neuroinflammation, and, critically, do not induce tolerance after chronic administration. These properties position Serodolin and MOA51 as highly promising next-generation analgesic candidates with clear translational potential.

## MATERIAL AND METHODS

### Compounds

Serodolin and MOA51 was synthetized by ICOA laboratory (Institut de Chimie Organique et Analytique, CNRS UMR 7311) as previously described ^21^.

### Radioligand binding competition assay

Radioligand binding studies were performed by Euroscreen Fast/ Epics Therapeutics S.A. Compounds have been prepared as 10 mM master solution in 100% DMSO. Serial dilutions were performed from master solution in 100% DMSO to obtain intermediate concentrations. CHO-K1 cell line was used during the assay. The radioligands used were [^3^H]-LSD for 5-HT_7A_ and 5-HT_6_ receptors, [^3^H]-8-OH-DPAT for 5-HT_1A_, [^3^H]-DOI for 5-HT_2A_ and 5-HT_2C_ and and [^3^H]-Spiperone for dopamine D_2L_ receptor. The reference ligands were 5-CT for 5-HT_7_R, serotonin for 5-HT_1A_R, 5-HT_2A_R, 5-HT_2C_R, Mianserin for 5-HT_6_R and Risperidone for D_2L_R. Compounds were tested by radioligand binding competition experiments at seven concentrations in duplicate: from 10 ^-12^M to 10^-5^M. For radioligand binding assay, competition binding was performed in duplicate in 96 wells plate (Master Block, Greiner, 786201) containing binding buffer, membrane extracts from CHO-K1 expressing the studied receptor, radiotracer and test compound. Nonspecific binding was determined by co-incubation with 200-fold excess of cold competitor. The samples were incubated in a final volume of 0,1 mL at a temperature and for a duration optimized for each receptor and then filtered over filter plates. Filters were washed six times with 0,5 mL of ice-cold washing buffer and 50 µL of Microscint 20 (Packard) are added in each well. The plates were incubated 15 min on an orbital shaker and then counted with a TopCountTM for 1 min/well. On each day of experimentation and prior to the testing of compounds, reference compounds were tested at several concentrations in duplicate to obtain a dose-response curve and an estimated IC_50_ values. Reference values thus obtained for the test were compared to historical values obtained from the same receptor and used to validate the experimental session. A session was considered as valid only if the reference value was found to be within a 0,5 logs interval from the historical value. For replicate determinations, the maximum variability tolerated in the test will be of +/-20% around the average of the replicates.

### Animals

C57BL/6 (Janvier and Charles River, France) or WT and 5-HT_7_R knockout (5-HT_7_R KO) mice maintained on a C57BL/6 background were used ^54^ (JAX Stock number 019453). For experiments, male and female animals (8 to 14 weeks old) were housed in a temperature (20-24°C)- and relative humidity 45 to 65%-controlled room and acclimated to a bright cycle (12/12h). All animal protocols were carried out accordingly with the French Government animal experiment regulations and were approved the Animal Ethical Committees (Comité d’Ethique pour l’Expérimentation Animale Orléans -CE03 and Auvergne -C2E2A) and accredited by the French Ministry of Education and Research (MESR) under national authorisation number C 45-234-12 (CBM-CNRS) and I-45-234-6 (TAAM). Animals used in this study will be treated accordingly to the guidelines of the Committee for Research and Ethical Issue of the I.A.S.P. (1983) and the European guidelines 2010/63/UE.

### Acetic acid writhing test in mice

The experiment involved 21 groups of 10 Swiss mice (Janvier Breeding Centre, France). Ten minutes before the intraperitoneal (i.p.) injection of a 1% acetic acid solution, mice were treated with different doses of Serodolin (0,1; 1; or 10 mg/kg) or MOA51 (0,1; 1; or 10 mg/kg) administered either orally or subcutaneously. The vehicle was 20% DMSO and 5% Tween^80^ in water for oral administration, or in saline for subcutaneous administration. Five minutes after the acetic acid injection, the number of writhing responses was recorded individually for 10 minutes.

### Toxicity studies

The experiment was performed on 7-week old mice male C57BL/6J mice (Janvier, France). Serodolin (10 mg/kg), MOA51 (1 mg/kg) or vehicle (20% DMSO, 5% Tween^80^ diluted in NaCl 0,9%) were administrated orally once daily during 10 consecutive days. Twenty-four hours after the first administration, a few drops of blood were collected from the submandibular (facial) vein and put in Multivette® 600 Lithium heparin LH tubes (Sarstedt, 15.1673). Haematological parameters were determined by Artimmune SAS (Orléans, France) using a multiparametric automated analyser (Olympus AU 400) based on photometric determination of analytes. On day 10, mice were sacrificed, and blood was collected intracardially in Multivette® 600 Lithium heparin LH tubes (Sarstedt, 15.1673). Hematological analysis was again determined by Artimmune SAS using the Olympus AU 400 analyser.

### Von frey test

The experiment was performed on 7-week old mice male C57BL/6J mice (Janvier, France). Mechanical allodynia was assessed using the Dynamic von Frey apparatus (Ugo Basile, Aniphy, Cat. No. 37450). Before to starting Von Frey measures, each mouse was placed in a transparent plexiglass box on an elevated grid platform during 20 minutes habituation period. On day 1, before inflammation was induced, baseline mechanical sensibility was determined by applying Von Frey filament to the plantar surface of left hind paw. Peripheral inflammation was induced by intraplantar injection of 10 µl of complete Freund’s adjuvant (F5881, Sigma) into the left hind paw. On day 2, mechanical allodynia was measured again to evaluate the effect of CFA-induced inflammation. Mice were divided into four groups. The first group consisted of vehicle-treated animals, the second group received CFA treatment only, the third group was treated with Serodolin (5 mg/kg, s.c.) or MOA51 (0,5 mg/kg, s.c.), and the fourth group received both CFA and subsequently Serodolin or MOA51 treatment. On day 3, the effects of the ligand on mechanical allodynia were evaluated using Von Frey test. The results were reported as a percentage of each animal’s baseline mechanical paw withdrawal threshold (set at 100%).

### Formalin induced inflammatory pain model in rats

This study was performed by ANS Biotech (Riom, France) on Sprague-Dawley male rats. A single subcutaneous administration of the Vehicle (20 % DMSO, 5 % Tween diluted in NaCL 0,9 %), Serodolin (10 mg/kg), MOA51 (1 mg/kg) or Morphine (4 mg/kg) was done 30 min before formalin injection on testing day (D0). On that day, rats received a unilateral injection of 50 µl of 2.5 % formalin solution into the plantar surface of the hind paw Hind paw licking time was recorded in consecutive 5-minute periods from 0 to 5 minutes (early phase) and 17 to 27 minutes (late phase) after the formalin injection. Immediately after behavioral test, dorsal root ganglion (DRG) and cerebrospinal fluid (50μl) were collected from all animals.

### Neuropathic allodynia cuff mice model

The experiment was performed on 7 weeks old mice male C57BL/6J mice (Charles River, France). Before surgery, each animal was tested with Von Frey calibrated filaments (Aniphy-Vivo Tech) to determine the basal sensibility level. The calibrated filaments were applied perpendicularly to the hindpaw according to a series of increasing forces ranging from 0,6 to 8 g. The pressure was stopped as soon as the filament bends. The mechanical threshold for removing the paw was determined by the first filament causing at least 3 removals of the paw over 5 applications. This test is carried out for two weeks before any surgery (baseline), then one week after surgery (in order to check the establishment of allodynia) then on the day of the time course: before administration of the compound and 30 min, 1h and 2h after injection. The neuropathic pain model used was induced by compression of the main branch of the right sciatic nerve. The animals were anesthetized with an intraperitoneal injection of 10 mL/kg of a mixture of zoletil (80 mg/kg) and xylazine (20 mg/kg) diluted in a 0.9% NaCl solution. The eyes were protected using Ocry-gel liquid eye gel (TVM Laboratory, Lempdes, France). After incision in the thigh, the common branch of the sciatic nerve was isolated then released using toothpicks previously sterilized (autoclave). The nerve was kept hydrated using a sterile physiological solution (NaCl 0.9%). A polyethylene sleeve (length 2 mm, internal diameter 0.38 mm, external diameter 1.09 mm; PE-20, Harvard Appartus, Les Ulis, France) was placed around the main branch of the right sciatic nerve. The incision was then sutured. Animals with surgery were part of the “Cuff” group. The “Sham” control animals were undergoing the same surgery without insertion of the cuff. The animals were placed back in their cage after surgery under a heat lamp; atipamezole (0,2 mg/kg, 5 mL/kg) was injected to reverse the anesthesia. Treatments with Vehicle (20 % DMSO, 5 % Tween diluted in NaCL 0,9 %), Duloxetin (30 mg/kg), as a reference molecule and different doses of Serodolin (10 mg/kg, 5 mg/kg, 1 mg/kg, 0,1 mg/kg) or MOA51 (1 mg/kg, 0,5 mg/kg, 0,1 mg/kg, 0,01 mg/kg) were started 15 days after the surgical procedure (n=7-8 mice/group). The effect of a first injection was monitored at different post-injection times (30 min, 1 hour and 2 hours) in “Cuff” animals presenting neuropathic allodynia and in “Sham” animals. 24 h after the last administration of molecules, animals were sacrificed and blood was collected to assess cytokines levels in plasma.

### Neuropathic allodynia spared nerve injury (sni) mice model

The experiment was performed on 8 weeks old male C57BL/6 mice (from Charles River). Mice were anesthetized and a complete ligature and transection of the common peroneal and tibial distal branches of the sciatic nerve was made leaving the sural branch intact. Mechanical threshold response of mice was measured with calibrated Von Frey filaments using the up/down method. Measures were performed as follow: one baseline measure before surgery, one measure at D+10, D+12, D+14, D+16 and D+18 before molecules administrations, and 1h, 2h post-drug administration. A supplementary measure 4h post-drug administration was performed at D+10 and D+18. Pregabalin (Tocris, ref 3775), the reference molecule was diluted at 0,3 mg/mL in PBS (Gibco, ref 14190-094). MOA51 and Serodolin were diluted in Vehicle solution (NaCl 0.9 % -DMSO 20 % -Tween80 5%). MOA51 was resuspended at 50 μg/mL and was administrated at a ratio of 100 μl per 10g (dose of 0,5 mg/kg). Serodolin was resuspended at 0,5 mg/mL and administrated at a ratio of 100 μl per 10g (dose of 5 mg/kg). Molecules were subcutaneously administrated (100 μl/10 g) for nine consecutive days, starting 10 days after surgery. At the end of the experience, transcardiac perfusion of 4 % Paraformaldehyde was performed in order to collect spinal cord for immunohistological analysis. Spinal cords were conserved 48h in sucrose (Thermo Scientific, 177140010) at 4°C and then embedding in a frozen mold with Tissue-Tek OCT (Optimal Cutting Temperature embedding medium, Cell Path, Newtown, UK, KMA-0100-00A).

### Preparation of samples and ENZYME-LINKED IMMUNOSORBENT ASSAY (ELISA)

Frozen Dorsal Root Ganglia L4-L5-L6 from “Formalin” experience, were resuspended in 150 µl of lysis buffer (TRIS 1M pH7,5 (Euromedex,200923-A), NaCl 5M (Euromedex, 1112-A), Sucrose (Thermo Scientific, 177140010), EDTA 0,5M pH8 (Euromedex, EU0084-B), SDS 10% (Biorad, 1610416) diluted in water added with PMSF, Na3VO4, Protease inhibitor (ChemCruz sc-24948-A) and tablets of PhosSTOP (Roche, 04906837001)) before sonication (Bioblock Sientific, Vibracell, 75115; amplitude 20%, pulses ON 5 sec OFF 3 sec). Then, IL-1β (R&D Systems, DY501-05), IL-10 (R&D Systems, DY504), TNF-α (R&D Systems, DY410) enzyme-linked immunosorbent assays were performed according manufacturer’s recommendations for cerebrospinal fluid and Dorsal Root Ganglia samples. Capture Antibody was diluted to the working concentration in PBS without carrier protein, added to the plates and incubated overnight at room temperature. Plates were washed three times with Wash Buffer (0,05 % Tween® 30 in PBS, pH 7,2-7,4). Plates were blocked by adding Reagent Diluent (R&D Systems, Catalog DY995) to each well. Plates were washed three times with Wash Buffer. 2X diluted in Reagent Diluent Dorsal Root Ganglia samples, pure cerebrospinal fluid samples and diluted standards in reagent diluent were added to the plates and incubated 2h at room temperature. Plates were washed three times with Wash Buffer. Detection antibody were diluted in Reagent Diluent and added to the plates for an incubation of 2h at room temperature. Plates were washed three times with Wash Buffer. Working dilution of streptavidin-HRP was added to the plate and incubated 20 min at room temperature in darkness. Plates were washed three times with Wash Buffer. Substrate solution (R&D Systems, Catalog DY999) was added to the plates and incubated for 20 min at room temperature in darkness. Stop solution (R&D Systems, Catalog DY994) was then added to the well to stop the reaction of the substrate. Optical density of each well was immediately determined using a microplate reader (CLARIOstar Plus, BMG Labtech) at two wavelengths (450 nm and 570 nm).

### Procartaplex ^tm^ assay

Using the manufacturer’s recommendation of The ProcartaPlex^TM^ Mouse and Rat Mix & Match Panels (Thermofisher, PPX-10-MXDJZNT), produced cytokines were detected in pure plasma blood sample from “cuff mice model” experiment. Before starting the assay, reagents were prepared (1X Wash Buffer, 1X Universal Assay Buffer and Standard Mix for generation of standard curve. Capture bead Mix was added to the plate and then beads were washed using a Hand-Held Magnetic Plate Washer. Pure plasma samples and standards were added to the plate and shacked at 600 rpm during the 2h of incubation at room temperature. Plate was washed three times with 1X Wash Buffer using a Hand-Held Magnetic Plate Washer. Biotinylated detection Antibody Mix was added to the plate and shacked at 600 rpm during the 30 min of incubation at room temperature. Plate was washed again three times with 1X Wash Buffer using a Hand-Held Magnetic Plate Washer. Streptavidin-PE was added to the plate and shacked at 600 rpm during the 30 min of incubation at room temperature. Plate was washed again three times with 1X Wash Buffer using a Hand-Held Magnetic Plate Washer. Reading Buffer was added to the plate and shacked at 600 rpm during the 5 min of incubation at room temperature. Plate was analysed on a xMAP^TM^ instrument and then final analysis were done on ThermoFisher website.

### Histology and immunofluorescence staining

After embedding in VWR®, OCT Embedding matrix for frozen sections (Catalog # 361603E), spinal cords from SNI mice model were sliced using a rotary cryostat (CM3050 S, Leica) at 10 μm thickness. Sections corresponding to spinal levels L3-L5, according to mouse atlas by Watson, Paxinos and Kayalioglu were used for immunofluorescence staining of Glial Fibrillary Acidic Protein (GFAP) for astrocyte detections, Iba-1 (Ionized calcium-binding adaptor molecule 1) for microglial detection and c-Fos as a marker of neuronal activity. Antigens recovering was performed by heat-induced technique with citrate buffer (citric acid- C0759, Sigma Aldrich; Sodium citrate- 1126, Euromedex) during 30 min at 80-90°C. Slides in hot citrate buffer were then placed on the bench at room temperature during 30 min to be cooled, followed by a brief rinse in distilled water and three washes of 5-minutes in 1X Tris Buffered Saline (TBS1X) (Euromedex, ET220-B). Slices on the slides were surrounded with Dakopen (Sigma, Z377821-1EA). Slides were incubated during 45 min at room temperature with the blocking solution (TBS 1X, 1 % BSA (Bovine Serum Albumin Fraction V pack, Sigma Aldrich, 810533), 10 % Foetal Bovine Serum, 0,3 % Triton (Biosolve, 201823) to block non-specific binding). Primary antibodies, anti-GFAP (Sigma, G6171), anti-Iba-1 (Abcam,5076) or anti-c-Fos (Proteintech, 66590-1-Ig) were applied overnight at 4°C in the blocking solution in darkness in the immunostaining moisture chamber. After three washes of 5 min with 1XTBS, slides were incubated during 1 hour in the dark with the appropriate secondary antibodies: Goat anti-mouse A488 (Invitrogen, A21121) for GFAP and c-Fos and Donkey-anti-Goat Alexa Fluor 555 (Thermofisher, A11055) for Iba-1. After an additional three -minutes 1X TBS washes, slides were mounted using Fluoromount-G™, with DAPI (Invitrogen, 00-4969-52) for nuclear staining. Fluorescence images were acquired using a videomicroscope (Axio Observer Z7, Carl Zeiss. SA). Quantification of GFAP, Iba-1 and c-Fos immunoreactivity was performed using ImageJ software.

### Stastistical analysis

All results are shown as mean ± SEM. For *in vivo* experiments, statistical analysis was performed using nonparametric Kruskal-Wallis test followed by Dunn post-test or a two-way ANOVA with Tukey post hoc test.

#### Cuff mice model

Graphs representing times courses or areas under the curve (AUC) were produced using GraphPad Prism 9 software. Results were expressed as group mean ± standard error of the mean (SEM). The AUCs were calculated using the trapezoid method, taking the value 0 as a reference. The analysis of the results of the von Frey tests was carried out using Statistica 13 software (Statsoft, USA). Statistical analysis was performed using multifactorial analysis of variance (ANOVA) on repeated measurements. Surgical procedures (“Sham” or “Cuff”) and different treatments were considered as intergroup factors. Measurement time was taken as a within-subject factor. Duncan’s test was used for post hoc comparison. The statistics are represented in the results section in relation to the “Cuff Vehicle” group with *p<0.05, **p<0.01 and ***p<0.001.

#### SNI mice model

Statistical analysis was performed using SigmaPlot 12.5 software. Two-way RM ANOVA (followed by Bonferroni post-hoc test) was used to analysed time course response at D+10 and D+18. Area under the curve was determined with the 1h and 2h post-drug administration measures to investigated tolerance of repeated administration of compounds.

## Author Contributions

**Fahima Madouri:** Writing-original draft, Conceptualization, Methodology and Investigation, Validation, Data curation**. Cyril Guimpied:** Methodology and Investigation, Data curation. **Enora Pigeon:** Methodology and Investigation, Data curation. **Nadège Hervouet-Coste:** Methodology and Investigation, validation. **David Gosset:** Methodology and Investigation. **Pascal Auzou:** Validation. **Mélanie Kremer:** Methodology and Investigation, Data curation, validation. **Quentin Leboulleux:** Methodology and Investigation, Data curation. **Marie-Aude Hiebel:** Conceptualization, Resources, validation. **Julie Le Bescont:** Methodology and Investigation, Resources, validation. **Gérald Guillaumet:** Conceptualization, Resources, validation. **Franck Suzenet:** Conceptualization, Resources, validation. **Patrick Baril:** Writing-review and editing, validation. **Raphaël Serreau:** Validation. **Séverine Morisset-Lopez:** Writing-original draft, Conceptualization, Validation, Funding -acquisition and Supervision.

## Funding Sources

This work was supported by C-VaLo, Région Centre Val de Loire, Technopole d’Orléans, La Ligue contre le cancer (IP/IQ-18220) and the Fédération de Recherche Physique et Chimie du vivant FR2708 Centre de Biophysique Moléculaire -Institut de Chimie Organique et Analytique.

## AKNOWLEDGMENTS

We thank Matthieu Refregiers for his guidance in immunofluorescence staining quantification and for insightful scientific discussions. We thank P@cific, the MO2VING platform for helpful discussions on immunofluorescence staining. We are grateful to Artimmune SAS for hematological studies, EuroscreenFast for radioligand binding studies, ANS Biotech for formalin rat model experiment, Compt Opt for cuff mice model and Phenotype expertise for SNI mice model. Some schematics were created with BioRender.com and used under an academic individual License.

## ABBREVIATIONS

5-HT_7_R: 5-HT_7_ receptor
GPCRs: G protein-coupled receptors
CNS: Central Nervous System
WT: Wild Type
5-HT_7_R KO mice: 5-HT_7_R knock out mice
DRG: Dorsal Root Ganglion
SNI: Spared Nerve Injury
GFAP: Glial Fibrillary Acidic Protein
Iba-1: Ionized calcium-binding adaptor molecule 1

## Notes

### Competing Interest Statement

The authors have declared no competing interest.

