## Supplemental figures for "TARGETING 5-HT7 RECEPTOR WITH BIASED LIGANDS TO ALLEVIATE PAIN AND SPINAL NEUROINFLAMMATION"

Madouri et al.

**Figure S1:** Serodolin and MOA51 *per.os administrations* do not induce toxicity or blood inflammation *in vivo*

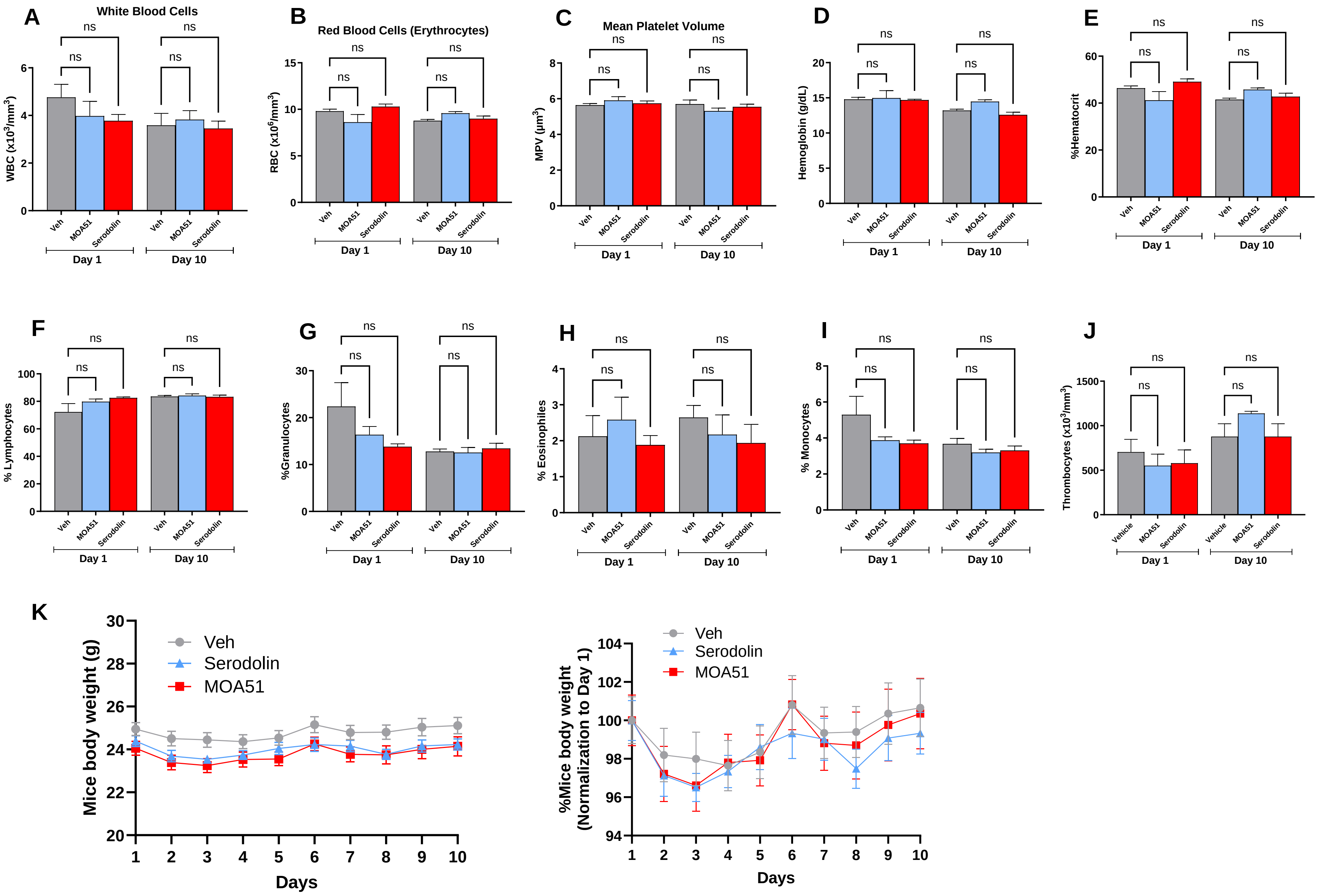

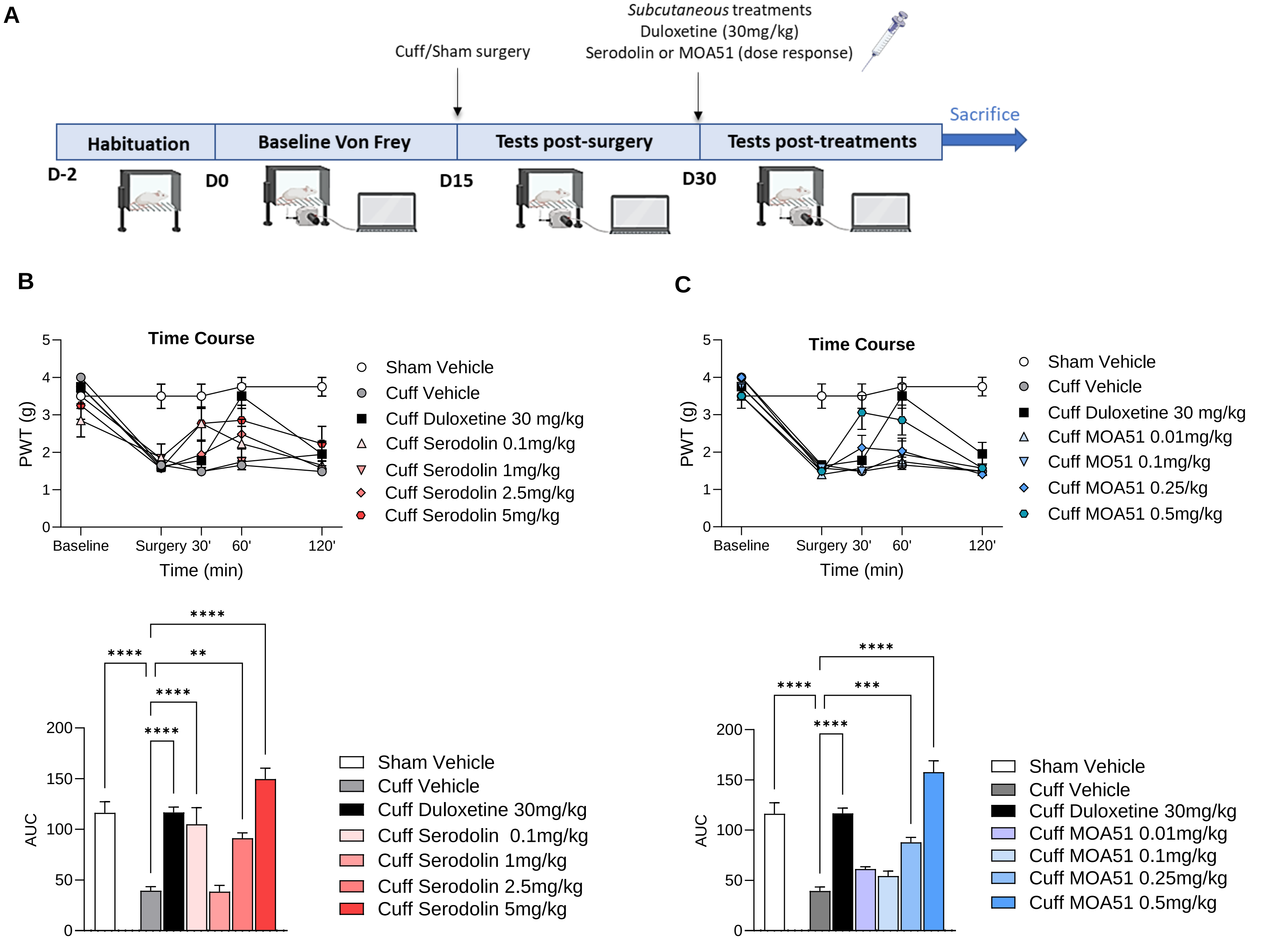


**Figure S2**: Serodolin and MOA51 subcutaneous administration have a complete and satisfactory effect on mechanical allodynia in the so-called "Cuff" model of sciatic nerve injury.

### SUPPLEMENTARY FIGURE LEGENDS

**Fig. S1. Oral administration of Serodolin and MOA51 does not induce toxicity or blood inflammation *in vivo*.** Acute (1 day) and chronic (10 days) oral administration of vehicle (20% DMSO, 5% Tween diluted in NaCl 0, 9%), Serodolin (10 mg/kg) or MOA51 (1 mg/kg) were performed. 24h after the first administration of compounds, a few drops of blood were collected from the submandibular (facial) vein (Day 1). For chronic administration, at day 10, blood was collected intracardially. Hematology was determined using multiparametric automate (Olympus AU 400) based on photometric determination of analytes: White Blood Cells (WBC) **(A)**, Red Blood Cells (RBC) **(B)**, Mean Platelet Volume (MPV) **(C)**, Hemoglobin **(D)**, Hematocrit percentage **(E)**, Percentage of Lymphocytes **(F)**, Granulocytes **(G)**, Eosinophils **(H)**, Monocytes **(I)** and Thrombocytes **(J)**. Mice body weight was measured daily during 10 days and represented in gramme and in a normalization to Day 1 **(K)**. Data are means ± SEM of values (n = 8 mice per group). ****P < 0.0001, ***P < 0.001, **P < 0.01, *P < 0.05. Statistical analysis was done using the Kruskal–Wallis test.

**Fig S2. Serodolin and MOA51 subcutaneous administration have a complete and satisfactory effect on mechanical allodynia in the so-called “Cuff” model of sciatic nerve injury. (A)** Design of “Cuff” model of sciatic nerve injury mice (n=7 mice per group). After a habituation period in the apparatus, baseline Von Frey measurements were realized on day 0. On day 15, cuff/sham surgery was performed and Von Frey measurement were realized to verify the effect of the surgery. Mice were treated with Duloxetine (30 mg/kg, s.c) or different doses of Serodolin subcutaneous administration (0,1 mg/kg; 1mg/kg; 2,5 mg/kg; 5 mg/kg) or MOA51 subcutaneous administration (0,01 mg/kg; 0,1 mg/kg; 0,25 mg/kg; 0,5 mg/kg). **(B)** The effect of Serodolin was evaluated on mechanical allodynia with Von Frey. **(C)** The effect of MOA51 was evaluated on mechanical allodynia with Von Frey. Results were represented as curve with Time Course and as Area Under the Curve (AUC). The results are expressed as the group means ± standard error of the mean (SEM). The AUCs were calculated using the trapezoid method taking the value 0 as a reference. The analysis of the results was carried out using Statistica 13 software (Statsoft, USA). A statistical analysis was performed using a multifactorial analysis of variance (ANOVA) during repeated measurements. The surgical procedures (“Sham” or “Cuff”) and the different treatments were selected as between-group factors. The measurement time was taken as a within-subject factor. The Duncan test was used for post-hoc comparison. The statistics are represented in the results section compared to the “Cuff Vehicle” group with *p<0.05, **p<0.01 and ***p<0.001.
